# DNA-based detection of *Aphanomyces cochlioides* in soil and sugar beet plants

**DOI:** 10.1101/2022.04.25.489453

**Authors:** Jacob R. Botkin, Cory D. Hirsch, Frank N. Martin, Ashok K. Chanda

**Affiliations:** Department of Plant Pathology, University of Minnesota, St. Paul, MN, 55108, U.S.A; USDA-ARS, Agricultural Research Service, Salinas, CA, 93905, U.S.A; University of Minnesota Northwest Research and Outreach Center, Crookston, MN 56716, U.S.A.

**Keywords:** *Aphanomyces cochlioides*, qPCR, detection, sugar beet, root rot

## Abstract

*Aphanomyces cochlioides*, the causal agent of seedling damping-off and Aphanomyces root rot (ARR) of sugar beet, causes yield losses in major sugar beet growing regions. Currently, a 4-week soil bioassay and a 2-day culture-based assay are used to diagnose presence of *A. cochlioides.* However, these assays can be time-consuming and lack sensitivity. In this study we developed a sensitive, specific, and rapid assay to detect and quantify DNA of *A. cochlioides*. We developed a TaqMan qPCR assay targeting a region of the mitochondrial genome of *A. cochlioides* representing a unique gene order for *Aphanomyces* with genus-specific primers and a species-specific probe. The qPCR assay detected *A. cochlioides* in 12 naturally infested field soil samples with disease severity index (DSI) values of 48-100, in sugar beet seedlings 5-7 days after planting, and with as little as 1 fg of pure *A. cochlioides* DNA. Adult sugar beet roots with ARR symptoms were sampled to further validate this qPCR assay. *Aphanomyces cochlioides* was detected in 95% of these samples using this qPCR assay, while only 23% of the same samples were positive using a culture-based assay. This shows the improved sensitivity of this qPCR assay for disease diagnosis and could provide growers with ARR risk of a field, which would help them make informed disease management decisions. However, further research is required to translate the results of this study to growers’ fields to quantify *A. cochlioides* with a high degree of accuracy.

## Introduction

Sugar beet (*Beta vulgaris* L.) is an economically important crop that is central to the multi-billion dollar sugar industry (Draycott, 2006). Sucrose extraction is performed on sugar beet taproots providing approximately 30% of the world’s sugar (Dohm et al., 2014). In 2021, 36.7 million tons of sugar beets were harvested in the United States with a market value of 1.8 billion dollars (USDA NASS, 2022). The Red River Valley (RRV) of Minnesota and North Dakota produces 51% of the total U.S. sugar beet crop. In the RRV, American Crystal Sugar Company (ACSC), a major sugar beet production cooperative, produces 3 billion pounds of sugar from 425,000 acres of sugar beet annually (ACSC, n.d.). Sugar beets are utilized as a fuel feedstock for ethanol production, and the pulp byproduct of sucrose extraction is used to produce non-starch livestock feed (Alexiades et al., 2018). Overall, sugar beets are agriculturally and economically valuable in the USA, as well as many other countries, including France, Canada, Russia, Ukraine, Turkey, Chile, Germany, Spain, Hungary, Sweden, the Netherlands, Australia, and Japan (Draycott, 2006; Harveson et al., 2007; Olsson et al., 2019).

The soil-dwelling oomycete, *Aphanomyces cochlioides*, is the causal agent of damping-off and root rot of sugar beet, causing yield losses in all major sugar beet growing regions. In the RRV, it was estimated that Aphanomyces root rot (ARR) occurs in half of all sugar beet fields, and in southern Minnesota it occurs in nearly all sugar beet fields (Beale et al., 2002; Harveson et al., 2007). The disease risk is higher in water-saturated regions of a field and affected area can range from a square meter to an entire field (Windels, 2000). Growing seasons with above average precipitation typically have a higher prevalence of ARR (Windels & Nabben-Schindler, 1996; Harveson, 2007). Severe ARR can reduce sucrose yields up to 27% and increase impurities in the roots which reduce its value for sucrose extraction (Olsson et al., 2011; Windels, 2000). ACSC estimates that its monetary losses are approximately 10 million dollars annually due to this disease (Harveson et al., 2007).

Aphanomyces seedling damping-off occurs after emergence, causing the hypocotyl to become necrotic and thread-like with the absence of foliar wilting. When ARR occurs on mature sugar beets the symptoms include chlorosis of older leaves, wilting during hot days, and necrotic water-soaked lesions that spread across the surface of the taproot, which result in stunting or plant death (Windels, 2000). Commercial sugar beet seeds are often treated with Tachigaren™ (hymexazol), which effectively protects from Aphanomyces and Pythium damping-off for a few weeks after planting. As sugar beets mature throughout the growing season, the chronic phase of ARR cannot be prevented with Tachigaren™. To reduce ARR severity, growers practice early planting of resistant sugar beet varieties, improve drainage by tiling and tillage practices, apply waste lime (precipitated calcium carbonate) to fields, implement a 3 to 5 year crop rotation with non-host crops, and control of alternative host weeds (Draycott, 2006; Harveson, 2007).

The hosts of *A. cochlioides* includes some Caryophyllales and dicots in the Amaranthaceae such as sugar beet (*Beta vulgaris* L.), table beet (*Beta vulgaris subsp. vulgaris* L.), spinach (*Spinacia oleracea)*, New Zealand spinach (*Tetragonia tetragonioides*), chard (*Beta vulgaris ssp*. *vulgaris*), cockscomb (*Celosia argentea* and *Celosia cristata* L.), and bouncing bet (*Saponaria officinalis*) (Wen et al., 2006; Windels, 2000). Additionally, wild weed and perennial flower species such as pigweed, lambsquarters (*Chenopodium album*), carpetweed (*Mollugo verticillata*), and fireweed can serve as inoculum reservoirs (Papavizas and Ayers, 1974; Grünwald et al., 2003; Harveson, 2007). ARR is an intractable disease due to its thick-walled oospores, which can persist in a dormant state for 10 to 20 years of adverse conditions without a host (Windels & Brantner, 2000; Harveson, 2007). Overwintering oospores serve as the primary inoculum for the following sugar beet crop (Harveson, 2007). When a host is present, under warm and saturated soil conditions, *A. cochlioides* oospores germinate by producing hyphae and form zoosporangia, which release large quantities of zoospores that migrate through soil water by sensing host root exudates and initiate infection (Islam, 2010). Overall, the ability to mass produce asexual zoospores that are capable of re-infecting host plants makes *A. cochlioides* a devastating sugar beet pathogen in warm and water-saturated soils.

Largely, traditional and molecular assays have been used to diagnose ARR, however these assays have limitations. A growth-chamber bioassay has been traditionally used to evaluate the ARR disease severity index (DSI) of infested field soil over a 4-week period (Fink and Buchholtz, 1954). However, this bioassay is limited by sensitivity and speed, and lacks the ability to quantify initial amounts of *A. cochlioides* inoculum in soil (Windels and Nabben-Schindler, 1996). Second, an ELISA targeting *Aphanomyces cochlioides* was developed to assess ARR of sugar beet. This assay is quantitative, but not specific to *A. cochlioides* and lacks sensitivity in samples with low inoculum amounts (Weiland & Shelver, 2004). A PCR assay was designed to amplify the *A. cochlioides* actin gene and the ITS region of the rRNA gene, but lacks specificity, and was only able to identify *A. cochlioides* with a restriction enzyme analysis of the amplicons (Weiland & Sundsbak, 2000). Another PCR assay specific to *A. cochlioides* was developed that multiplexed a primer pair specific to *A. cochlioides* with another primer pair specific to *A. euteiches* (Vandemark et al., 2000). This allowed the detection of both pathogens to be performed in one PCR run, but lacked the ability to quantify the pathogens. Recently, a specific qPCR assay was designed to detect and quantify *A. cochlioides* from infested soil using the internal transcribed spacer (ITS) region of the rRNA gene (Almquist et al., 2016). This qPCR assay had a reported limit of detection in the range of 1-50 oospores per gram of soil under artificial infestation depending on soil clay content. However, this qPCR assay did not detect *A. cochlioides* in 68.9% of naturally infested soil samples that were positive for *A. cochlioides* with the bioassay with a DSI below 80. Largely, traditional and molecular assays have been used to evaluate ARR, however these assays have apparent drawbacks.

The limitations of the available assays for ARR highlight the importance of designing an effective assay to detect and quantify *A. cochlioides* in soil and *in planta*. This study developed a qPCR assay that is sensitive, specific, and able to accurately quantify *A. cochlioides* in naturally infested field soil and infected plant samples. This qPCR assay will be useful for growers to accurately assess the Aphanomyces root rot risk level of a field prior to planting sugar beets, without having to wait 4 to 5 weeks for the soil bioassay results. This time is valuable because management decisions are determined by the infestation risk level in the field. Additionally, this qPCR assay could be useful for sugar beet diagnosticians to assess sugar beets infected with *A. cochlioides*. Overall, sugar beet growers have a need for a more sensitive, accurate and rapid assay that can consistently quantify initial infestation levels of naturally infested field soils. Having this information will help growers make informed decisions, such as planting date, selecting a resistant cultivar, use of appropriate dose of Tachigaren seed treatment, applying sugar beet factory waste lime, managing soil moisture, length of crop rotation, and other sanitation methods.

## Materials and Methods

### qPCR primer and probe development

To design the primers and probe, the mitochondrial genomes of 14 *Aphanomyces* isolates representing 8 species, including *A. cladogamus, A, cochlioides* and *A. euteiches*, were assembled and compared (F. Martin, unpublished). These included the mitochondrial genome of 5 *A. cochlioides* isolates, 13-35-5 (MN), 14-SMAN-M-1 (MN), 61ss (TX), 64ss (TX), C10 (ND), that were assembled from Illumina sequencing data. In addition, 250 available mitochondrial genomes from *Pythium, Phytophthora*, downy mildews and other oomycetes were used to identify differences in conserved gene order of *Aphanomyces* spp.. Within the mitochondrial genomes, 4 loci were specific to Saprolegniales, and 3 loci were specific to *Aphanomyces*. The forward primer (Aphcoc-F) was designed to anneal to a conserved region just downstream from the mitochondrial *rps4* gene for any *Aphanomyces* spp.. (Supplementary Figure S1). The reverse primer (Aphcoc-R) was designed to anneal to the 5’ end of the mitochondrial *nad2* gene for the plant pathogens *A. cladogamus*, *A. cochlioides*, and *A. euteiches*. The probe (Aphcoc-Pr) was designed to anneal to a region just before the 5’ end of the mitochondrial *nad2* gene of *A. cochlioides* alone. Using these primers for amplification of *A. cochlioides* resulted in a 234 bp amplicon.

### Development of qPCR detection assay

Quantitative PCR (qPCR) was executed using a LightCycler 480 Instrument II (Roche Life Science, Pleasanton, CA, USA) with a reaction volume of 25 µL per sample. The standard *A. cochlioides* assay reaction contents consisted of 1x PerfeCTa Multiplex qPCR ToughMix (Quantabio, Beverly, MA, USA), 0.6 µL (10 µM) forward primer (Aphcoc-F, 5’-GACCCTATTTAAAAATAGGTAT-3’), 0.6 µL (10 µM) reverse primer (Aphcoc-R, 5’-AAAAATTCAGGAATTAGAAATAAA-3’), 0.2 µL (10 µM) probe (Aphcoc-Pr, 5’ 6-FAM ATATAAATAATAATAATACATACATGATT-3’ BHQ), 1 µL template DNA, and the remaining volume made up to 25 µL with molecular grade water. The Roche LightCycler instrument measured the intensity of FAM (465-510 nm) for the *A. cochlioides* probe (Aphcoc-Pr). The thermal cycler conditions were as follows; a reaction initialization at 95°C for 3 minutes, followed by 40 cycles of denaturing at 95°C for 15 seconds and annealing at 60°C for 45 seconds, with a final elongation at 37°C for 30 minutes. Each qPCR plate included a no template negative control and a positive control consisting of 1 ng *A. cochlioides* DNA, which was also used in the standard curve. The standard curve was applied externally to each run using a fit point analysis.

A standard curve was made using eight 10-fold dilutions of *A. cochlioides* isolate 13-69-4 DNA, from 1 ng/µL to 0.1 fg/µL. The 1 ng/µL DNA dilution level was initially quantified using a Qubit fluorometer (Thermo Fisher Scientific, Waltham, MA, USA). The 0.1 ng concentration was prepared by adding 100 µL of the 1 ng/µL DNA dilution level to 900 µL molecular-grade water, then vortexing for 3 s, and repeating the process for all eight dilutions. Aliquots of 10 µL and 100 µL were prepared for each DNA dilution, and all DNA standards were stored at −80°C until use. One microliter of each DNA standard was tested with the qPCR assay to attain Ct values for the 1 ng to 1 fg dilutions. The standard curve Ct values were saved as an external standard curve, which was imported for each run using the fit point analysis.

The specificity and inclusivity of the qPCR assay was evaluated using the standard primer and probe set (Aphcoc-F, Aphcoc-R, Aphcoc-Pr). First, the specificity was evaluated *in silico* using NCBI’s nucleotide BLAST. Next, the qPCR assay was tested against DNA from *Beta vulgaris*, *A. euteiches*, *A. cladogamus*, Ascomycota fungi, non-Aphanomyces species of Oomycota, and Protozoa isolates (Table 1). Additionally, DNA from 14 *A. cochlioides* isolates obtained in the Red River Valley of Minnesota and North Dakota, southern Minnesota, and Texas were included to test the inclusivity of the qPCR assay (Table 1). *A. cochlioides* isolates were grown on 0.5X PDA and DNA was extracted from mycelial tissue using the FastDNA Spin Kit for Fungi (MP Biomedicals, Irvine, CA, USA) following the manufacturer’s specifications. All DNA samples were quantified using the Nanodrop Spectrometer (Thermo Fisher Scientific, Waltham, MA, USA) and 10 ng of DNA from each sample was used to test the specificity and inclusivity of the qPCR assay.

**Table 1.**
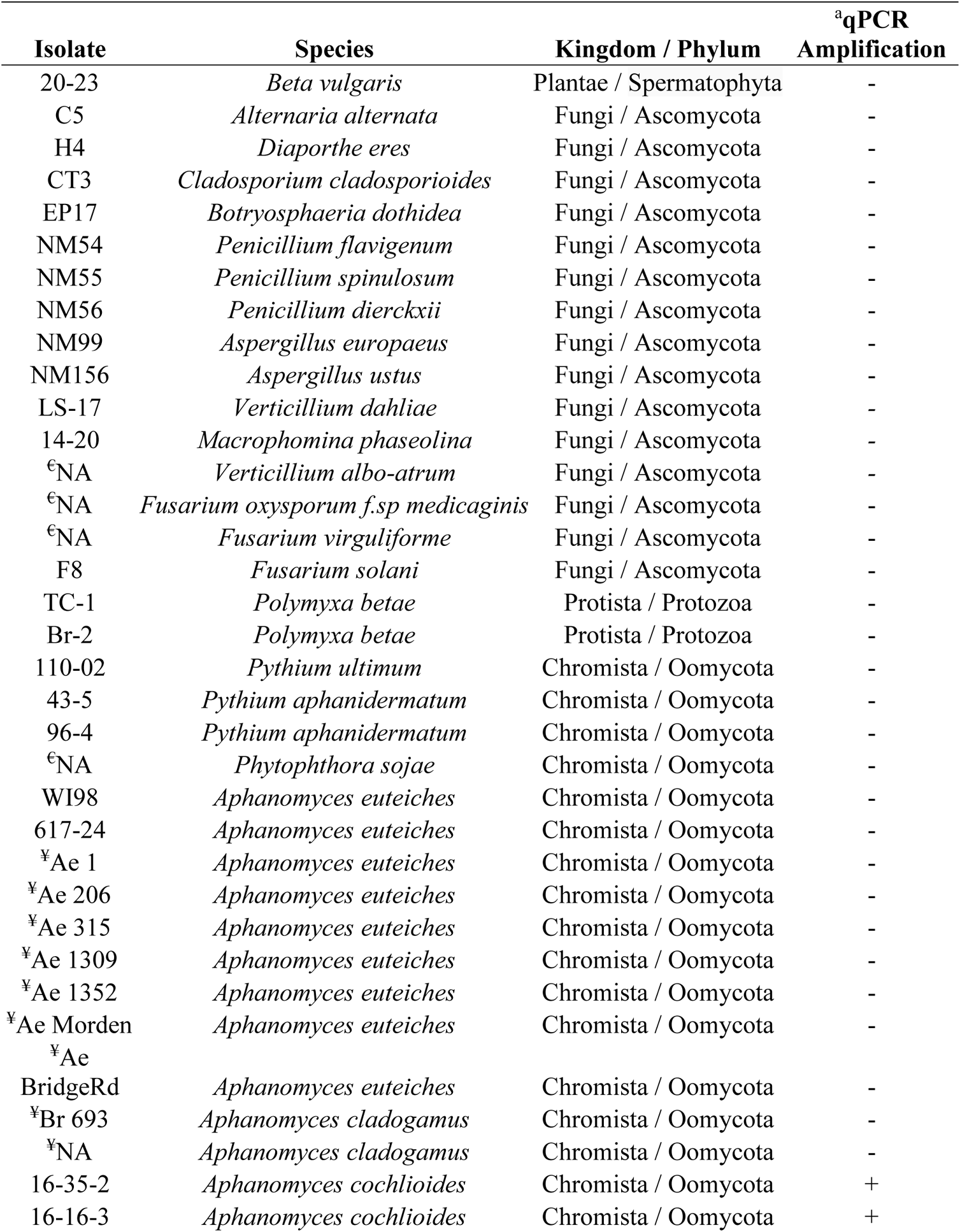
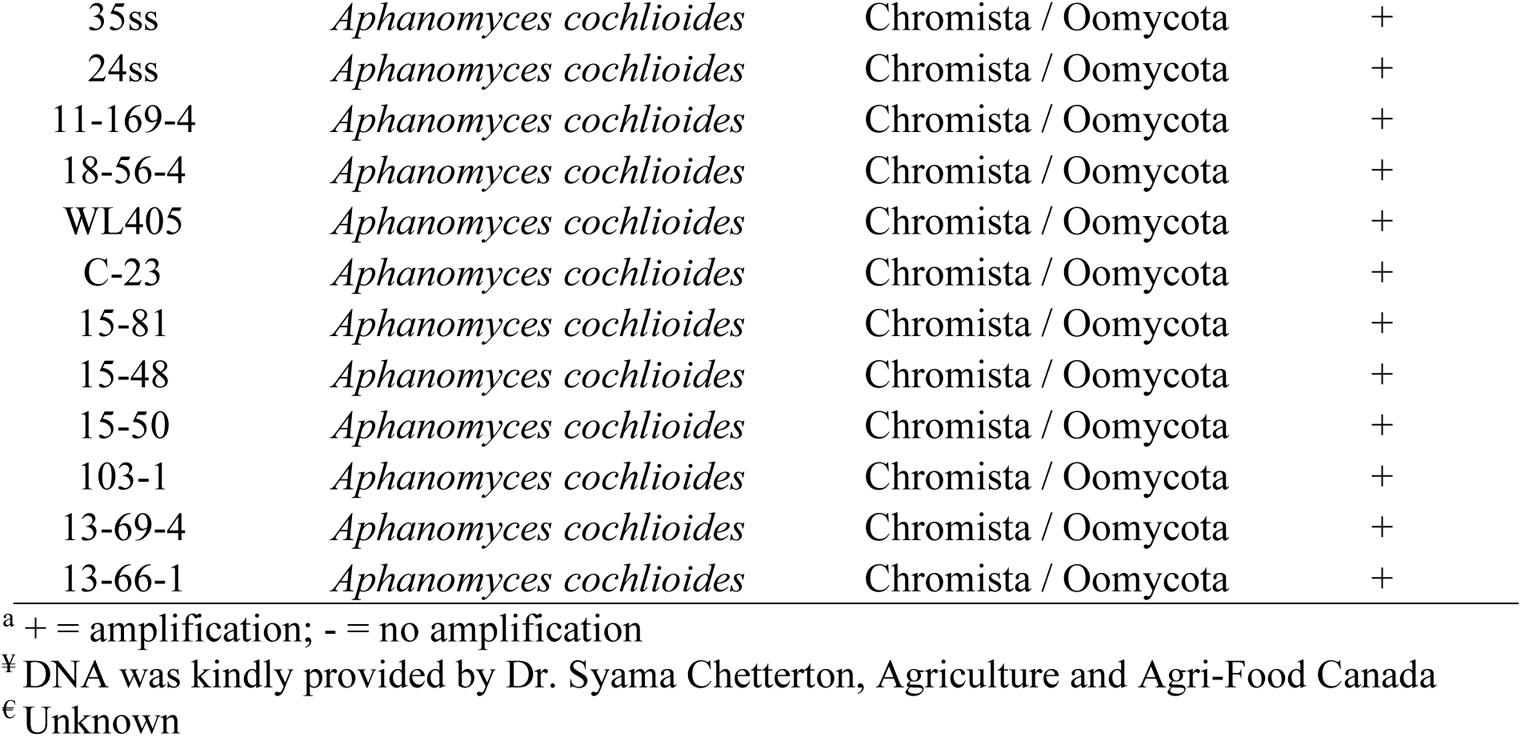
The isolates of fungi, oomycetes, protista and sugar beet used to test the specificity of the qPCR assay

### Naturally infested soil samples

Twelve naturally infested soil samples were collected from fields in the RRV of Minnesota and North Dakota, and Southern Minnesota from 2017 to 2020 (Table 2). Soils were used directly to conduct the bioassay and subsampled for DNA extraction. To prepare soil for subsampling, each soil was mixed thoroughly by hand in a 5-gallon bucket, 1 kg of soil was removed and homogenized with a mortar and pestle then passed through a 3 mm sieve. Homogenized soil samples were poured into a large tray and thoroughly mixed by hand. Three 500 mg and 5 g soil subsamples were removed and placed in 2 mL micro-centrifuge tubes and 50 mL centrifuge tubes, respectively, and stored at 4°C until DNA extraction.

**Table 2.** Location and year of collection for 12 soils naturally infested with *Aphanomyces cochlioides* in Minnesota and North Dakota.

| Sample Name | County | State | Year |
| --- | --- | --- | --- |
| AC10 S3 | Marshall | Minnesota | 2017 |
| AC10 S4 | Marshall | Minnesota | 2017 |
| 19-38 | Redwood | Minnesota | 2019 |
| 19-43 | McLeod | Minnesota | 2019 |
| 19-47 | Renville | Minnesota | 2019 |
| 19-59 | Renville | Minnesota | 2019 |
| 19-60 | McLeod | Minnesota | 2019 |
| SM-AN | Renville | Minnesota | 2019 |
| Climax South | Polk | Minnesota | 2020 |
| Grandin | Traill | North<br>Dakota | 2020 |
| Perley | Norman | Minnesota | 2020 |
| Climax North | Polk | Minnesota | 2020 |

### Aphanomyces root rot bioassay for infested soils

The ARR DSI of infested soils was assessed using the ARR growth chamber bioassay. For each soil sample, 25 seeds of Crystal 257 RR were planted in pots (10 x 10 x 10 cm) filled with naturally infested field soil. Each pot was considered a replicate and six replicates were used for each soil sample. Seed was commercially treated with a metalaxyl (0.15 g a.i. per kg seed) and thiram (2.49 g a.i. per kg seed) to protect against *Pythium* spp. Pots were arranged in a randomized block design and incubated in a growth chamber for 1 week at 20°C, followed by 3 weeks at 25°C, with a 14-hour photoperiod. To create favorable infection conditions for *A. cochlioides* pots were watered once per day. Seedlings were counted every day during emergence and three times weekly thereafter. Dying seedlings were removed, surface sterilized in a 0.5% sodium hypochlorite solution, rinsed with sterile deionized water, plated in sterile deionized ultra-filtered water (Fisher Chemical, Waltham, MA, USA) and microscopically evaluated to verify presence of *A. cochlioides* zoosporangia or other pathogens after 24 hrs. Four weeks after planting, remaining seedlings were removed from soil, washed, and rated using a 0 to 3 disease severity scale (Figure 1). The number of seedlings that died during the 4-week bioassay along with the disease severity ratings were used to calculate the DSI (Equation 1). The DSI values range from 0 to 100, with 0 representing no disease, and 100 representing the most severe ARR (Windels and Nabben-Schindler, 1996; Beale et al., 2002).

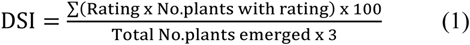

**Figure 1.**
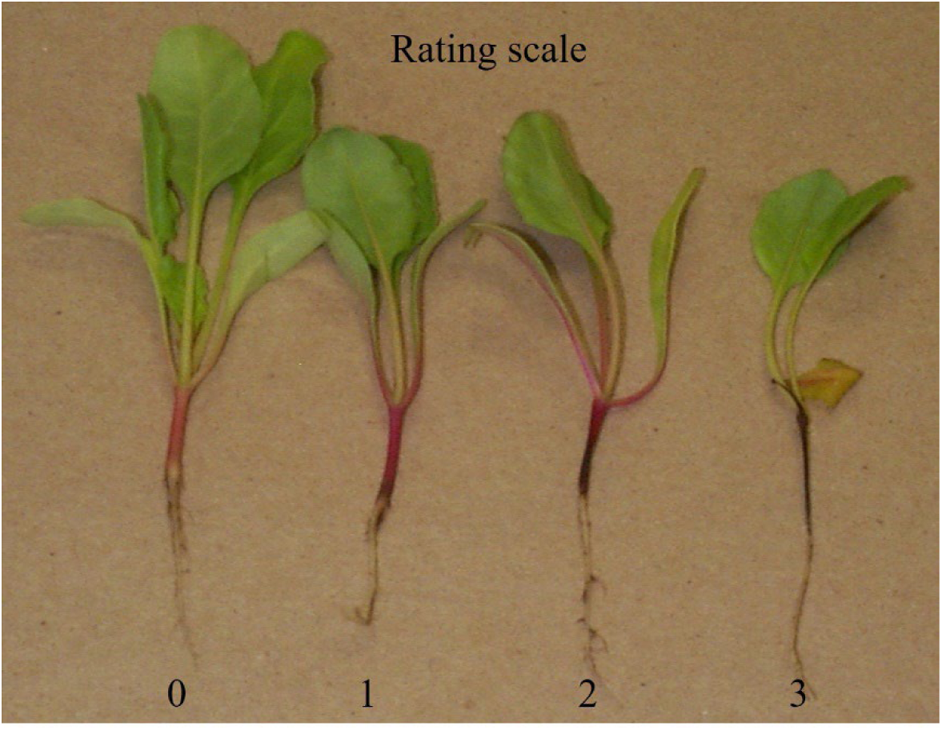
Disease severity rating scale used for the Aphanomyces root rot bioassay.

Equation 1. The Aphanomyces root rot disease severity index (DSI) is calculated based on the number of seedlings with a 0 to 3 rating (0 = healthy hypocotyl, 1 = light brown hypocotyl discoloration, 2 = moderate hypocotyl discoloration, 3 = severe hypocotyl discoloration or dead plant) (Windels & Nabben-Schindler, 1996).

### Naturally infected plant samples

Plant samples of diseased adult sugar beet roots were submitted to the sugar beet pathology lab at the University of Minnesota, Northwest Research and Outreach Center, Crookston for diagnosis during the summer of 2020. A total of 60 adult sugar beet roots were submitted from 11 fields in the RRV of Minnesota and North Dakota, and Southern Minnesota (Table 3). The soil was removed from roots with a brush under running water, an image was taken, and observations of root rot symptoms were recorded for each individual root. Of these samples, 40 roots with ARR symptoms such as irregular water-soaked or necrotic lesions, root constriction, scarring and malformation were selected to be tested for *A. cochlioides* using the culture-based and qPCR assay. Tissue subsamples, approximately 1 x 1 x 0.5 cm (150 mg), were excised from the margin of necrotic lesions on the root with a sterile scalpel. Two subsamples were taken right next to each other (Figure 2). One subsample was used for a culture-based assay and the other subsample was flash frozen in liquid nitrogen for DNA extraction and the qPCR detection assay.

**Figure 2.**
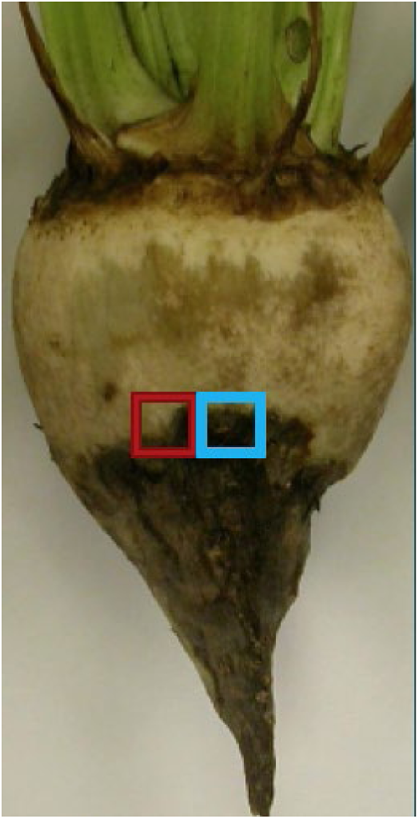
Locations of two subsamples taken from an adult sugar beet root for culture-based assay (blue square) and qPCR assay (red square).

**Table 3.**
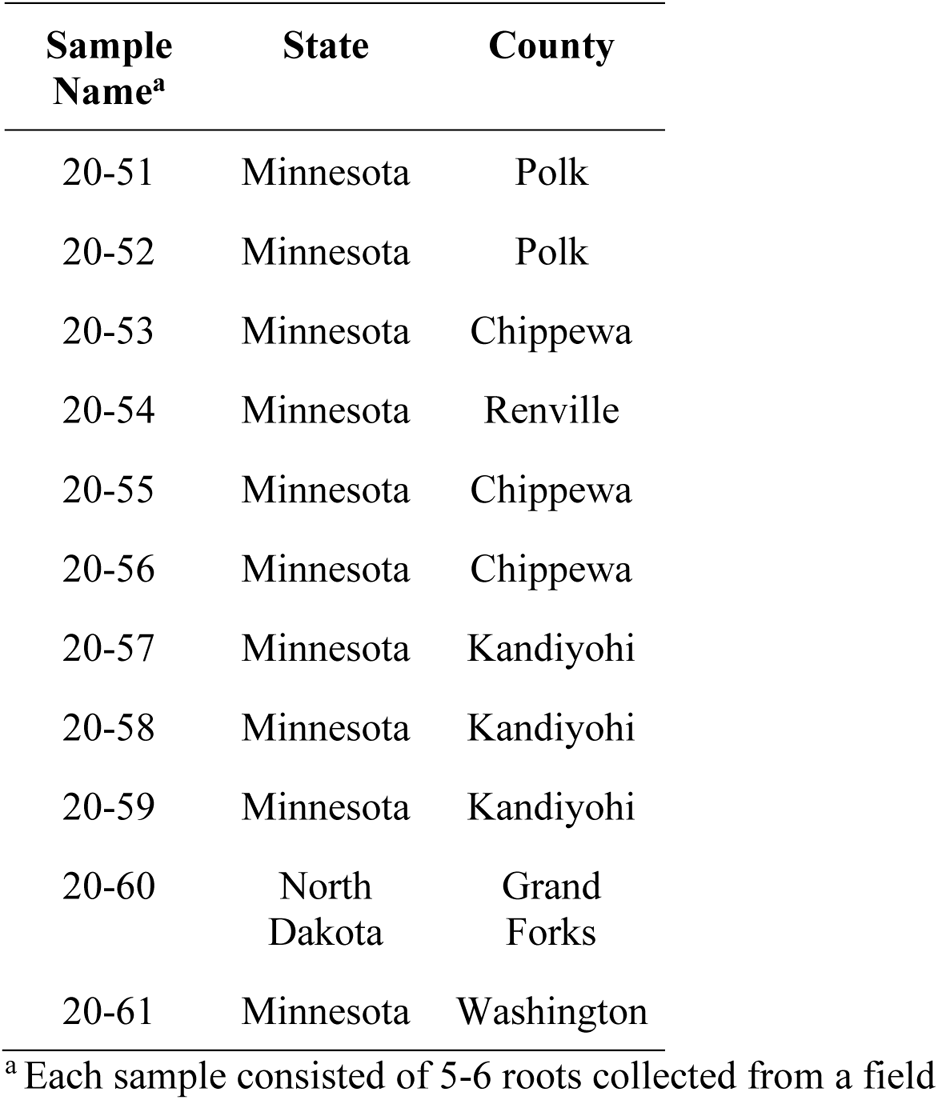
Location of sugar beet fields from which sugar beet roots naturally infected with *Aphanomyces cochlioides* were received during the summer of 2020.

### Culture-based assay for infected plants samples

To conduct the culture-based assay, excised tissue subsamples were surface sterilized as previously noted, rinsed with sterile deionized water, plated in 10 mL of sterile deionized ultra-filtered water, and incubated at room temperature. Water cultures of plant subsamples were examined with a stereo microscope one to two times a day over a three-day period. The presence of nonseptate hyphae, zoosporangia, zoospores or oospores was treated as confirmation that the infection was caused by *A. cochlioides*. Attempts were made to isolate *A. cochlioides* from diseased root tissue by culturing on PDA-RP [potato dextrose agar amended with rifampicin (18.8 mg/L) and penicillin G (50 mg/L)], or PDA-MBR [potato dextrose agar amended with metalaxyl (30 mg/L), benomyl (5 mg/L), and rifampicin (18.8 mg/L)] media.

### DNA extraction from soil and plant tissue

The FastDNA Spin Kit for Soil and Plants (MP Biomedicals, Irvine, CA, USA) was used for soil DNA extractions based on previous qPCR assays for soilborne plant pathogens (Almquist et al., 2016; Bilodeau et al., 2012; Wallenhammar et al., 2012). The standard protocol for the FastDNA Spin Kit for Soil and Plants was optimized for extracting *A. cochlioides* DNA by increasing the duration and speed of mechanical lysis. The optimized procedures for soil and plant tissue are described below.

For naturally infested soil samples, each sample was 500 mg (dry wt.), which was stored at 4°C until DNA extraction. An extra ceramic bead was added to each sample tube with Lysing Matrix E. Samples were subjected to mechanical cell lysis for 80 seconds using the FastPrep-24™ 5G (MP Biomedicals, Irvine, CA, USA) instrument at speed 7.5 m/s, or for 160 seconds with the Bead Beater instrument (BioSpec Products, Bartlesville, OK, USA) at the highest speed. The rest of the protocol was conducted according to the manufacturer’s recommendations. Final eluted soil DNA samples were stored at −20°C.

For naturally infected plant samples, tissue was flash frozen in liquid nitrogen and stored at −80°C. Each 150 mg (fresh wt.) tissue sample was placed in Lysing Matrix A, and an extra ceramic bead was added to each sample tube. Mechanical cell lysis was done for 80 seconds using the FastPrep instrument set at speed 7.5 m/s, or for 160 seconds with the Bead Beater instrument at the highest speed. The rest of the protocol was followed according to the manufacturer’s recommendations, and final eluted DNA samples were stored at −20°C.

### Detection of *A. cochlioides* in sugar beet seedlings

Four pots (10 cm x 10 cm x 10 cm) were filled with naturally infested field soil (Perley, MN, Table 2) and four pots (10 cm x 10 cm x 10 cm) were filled with the same soil, which had been recently used to conduct the bioassay (referred to as used bioassay soil from here on). Ten untreated sugar beet seeds (cultivar Crystal 257 RR) susceptible to *A. cochlioides,* were planted in each pot, and pots were placed in the growth chamber at 25°C with a 14-hour photoperiod. Seedlings were carefully uprooted at 3, 5, 7, 9, and 11 days after planting (DAP) to keep roots intact. Root tissue was washed under running water for 2 minutes on a screen, and a paintbrush was used to remove excess soil. Root infection was confirmed by examination with a stereo microscope. For each pot, all ten seedlings were removed, 5 seedlings were placed in a microcentrifuge tube, representing 2 replicates per collection time point. Sample tubes containing 5 seedlings were immediately frozen in liquid nitrogen and stored at −80°C until use. DNA extractions were performed using the FastDNA Spin Kit for Plants (MP Biomedicals, Irvine, CA, USA) using the optimized procedure described previously, and DNA was stored at −20°C until use. The qPCR assay was conducted for all seedling DNA samples using the procedure described previously, and Ct values were obtained using the fit point analysis.

### Detection of *A. cochlioides* in naturally infested soils

A total of 12 field soil samples naturally infested with *A. cochlioides* were used to validate the qPCR assay. All samples were subjected to the growth chamber bioassay for *A. cochlioides* described earlier. DNA was extracted from two samples received in 2017 using the DNeasy PowerMax Soil Kit starting with a 5 g soil sample and following the manufacturer’s recommendations. DNA was extracted from ten naturally infested field soil samples received in 2019 and 2020 using the FastDNA Spin Kit for Soil optimized protocol. All 12 samples were tested using the qPCR assay following the procedure described above. Ct values were obtained using the fit point analysis.

### Detection of *A. cochlioides* in naturally infected plant samples

Three *A. cochlioides* detection methods were used in parallel on diseased adult sugar beet roots collected from 11 fields known to have ARR in past growing seasons (Table 3). The detection methods included a visual assessment for ARR symptoms, the culture-based assay, and the qPCR assay. Adjacent subsamples, excised from each diseased adult sugar beet root, were either plated in sterile water cultures to recover the pathogen or subjected to DNA extraction using the FastDNA Spin kit for plants optimized protocol to obtain DNA, and the qPCR assay was used to confirm the presence of *A. cochlioides* DNA. Ct values were obtained using a fit point analysis and the results of the three detection methods were compared.

### Detection of *A. cochlioides* in artificially infested potting soil

The growth chamber soil bioassay and qPCR assay were performed on potting soil infested with different amounts of *A. cochlioides* isolate 103-1 oospores. Oospores were produced on excised sugar beet hypocotyls following the procedure described earlier (Dyer & Windels, 2003). Oospores were quantified with a Speirs-Levy counting chamber, added to autoclaved fine vermiculite, and air dried at 25°C for 2 days. Potting soil (Sun Gro Professional Growing Mix, Sun Gro Horticulture, Agawam, MA, USA) was amended with 60% sand (Difco Laboratories Inc, Franklin Lakes, NJ, USA) by mass and autoclaved. The oospore-vermiculite inoculum was weighed and mixed into the soil at 0, 1, 5, 10, 20, 50, and 100 oospores/g soil (dry wt.) in trays by hand. The infested soil was watered with reverse osmosis water, and air dried at 25°C for 10 days. Infested soil was plated on PDA-RP medium to confirm *A. cochlioides* mycelium was inactive. Five seeds from the susceptible sugar beet cultivar Crystal 093 RR were planted in infested soil following established methods, with eight technical replicates for each soil-oospore dilution (Windels & Brantner, 2001). After mixing oospores into soil, two 500 mg samples for each infestation level were removed and used for DNA extraction following the FastDNA Spin Kit for Soil optimized procedure. Finally, the qPCR assay and the ARR bioassay were conducted, as described above, to assess the Ct value and ARR DSI for each oospore infestation level.

## Results

### Specificity of the qPCR assay

NCBI’s nucleotide BLAST was used to computationally evaluate the specificity of the *A. cochlioides* forward (Aphcoc-F) and reverse (Aphcoc-R) primers, which were designed to be specific to the genus *Aphanomyces* and *Aphanomyces* spp. that are plant pathogens, respectively, as well as the probe (Aphcoc-Pr) that is specific to *A. cochlioides*. The results showed both the forward and reverse primers and probe have 100% identity with the *A. cochlioides* isolate 103-1 (accession JAHQRK010000050.1) and did not have high similarity with plant pathogens outside of the genus *Aphanomyces.* Additionally, the forward primer has 100% identity to a region in the mitochondrial genomes of *A. astaci* isolate NJM9701 (accession KX405005.1), *A. invadans* isolate AP03 (accession KX405005.1), and other *Aphanomyces* spp. Finally, the reverse primer has 100% identity to *Aphanomyces euteiches* strain ATCC 201684 (accession VJMJ01000379.1).

To evaluate the specificity of the qPCR assay, it was tested against 10 ng of DNA from a variety of other taxa, and multiple isolates of *A. cochlioides*. These included 9 isolates of *A. euteiches*, 2 isolates of *A. cladogamus*, 15 species of Ascomycota fungi, 3 non-Aphanomyces species of Oomycota, and a species of Protozoa (Table 1). DNA from 14 isolates of *A. cochlioides* originated from North Dakota, Minnesota, and Texas was used to test the inclusivity of the qPCR assay. The results of this analysis showed that the qPCR assay is specific to *A. cochlioides*, as DNA amplification did not occur for any species tested other than *A. cochlioides*. Also, the qPCR assay was shown to be inclusive, with the DNA from each *A. cochlioides* isolate being detected consistently (Table 1).

### Sensitivity of the qPCR assay

A standard curve of 10-fold serial dilutions of pure *A. cochlioides* genomic DNA was tested with the qPCR assay in order to assess the linearity of amplification and limit of detection (LOD). The LOD of the qPCR assay was 1 fg, which had a mean Ct value of 34.66 (Table 4). Each standard from 1 ng to 1 fg was detected in every technical replicate. For DNA standards that were detected consistently, the difference between mean Ct values of concentrations ranged from 3.41-4.15, giving an amplification efficiency of 83.3% (Figure 3). However, since the standards were not tested in the background of soil or sugar beet DNA, the qPCR amplification efficiency and the LOD may differ in DNA samples of naturally infested soils or infected plants that may contain different levels of PCR inhibitors.

**Figure 3.**
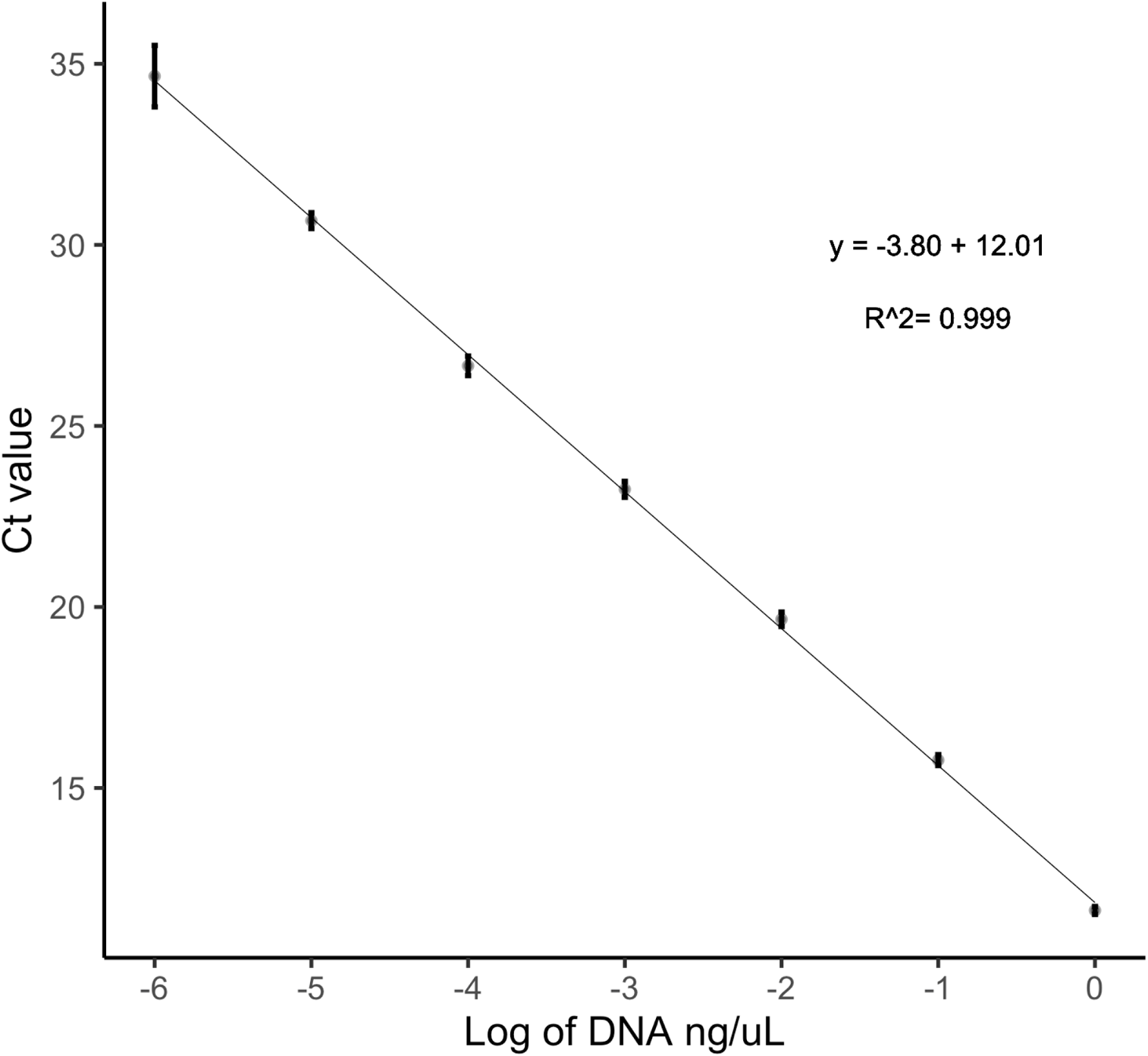
Standard curve of 10-fold dilutions of pure *Aphanomyces cochlioides* isolate 13-69-4 genomic DNA standards. Standard error bars are displayed with vertical black lines.

**Table 4.**
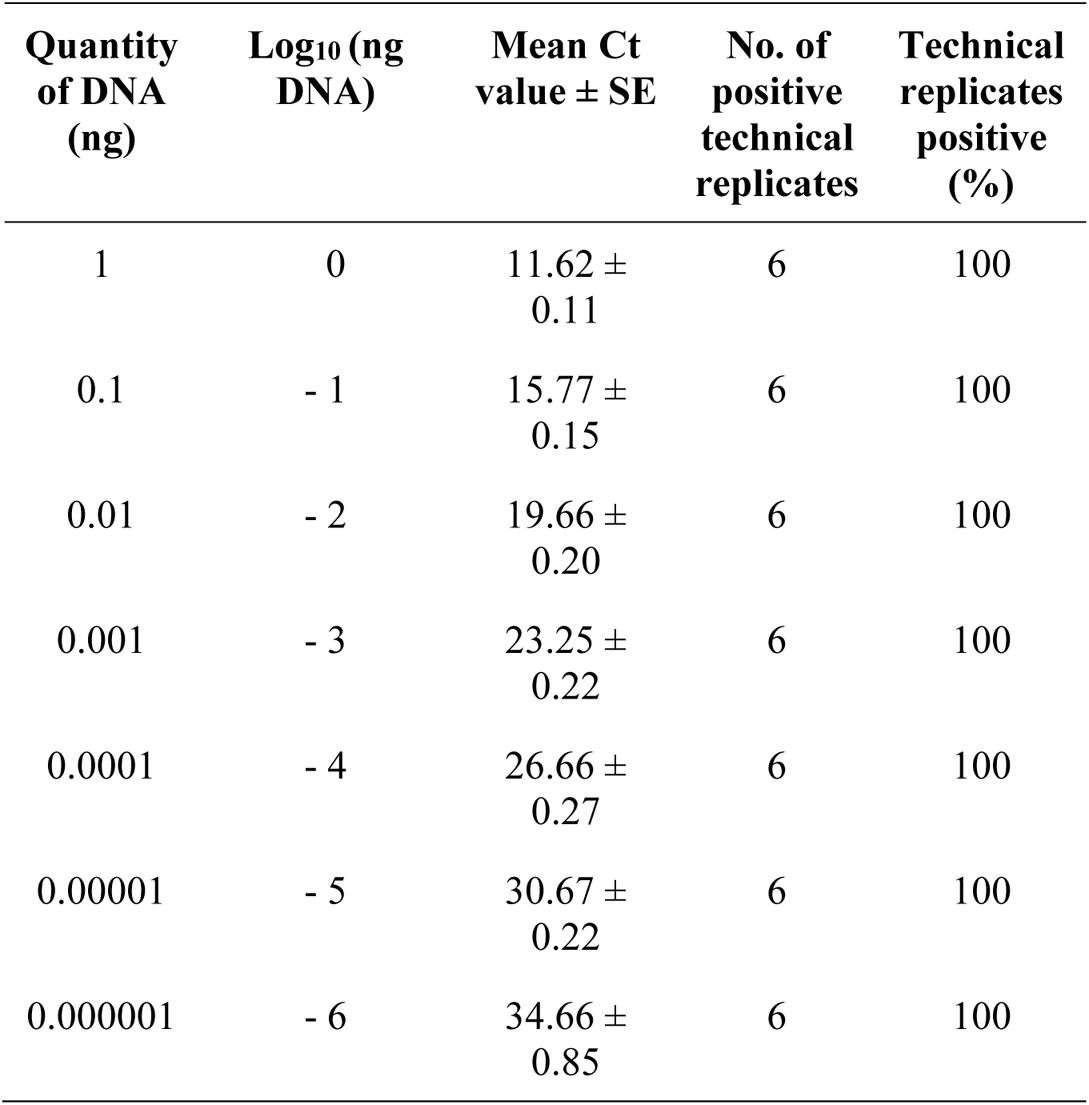
Standard curve of 10-fold serial dilutions of pure *Aphanomyces cochlioides* genomic DNA.

| Quantity<br>of DNA<br>(ng) | Log<sub>10</sub> (ng<br>DNA) | Mean Ct<br>value ± SE | No. of<br>positive<br>technical<br>replicates | Technical<br>replicates<br>positive<br>(%) |
| --- | --- | --- | --- | --- |
| 1 | 0 | 11.62 ±<br>0.11 | 6 | 100 |
| 0.1 | - 1 | 15.77 ±<br>0.15 | 6 | 100 |
| 0.01 | - 2 | 19.66 ±<br>0.20 | 6 | 100 |
| 0.001 | - 3 | 23.25 ±<br>0.22 | 6 | 100 |
| 0.0001 | - 4 | 26.66 ±<br>0.27 | 6 | 100 |
| 0.00001 | - 5 | 30.67 ±<br>0.22 | 6 | 100 |
| 0.000001 | - 6 | 34.66 ±<br>0.85 | 6 | 100 |

### Detection of *A. cochlioides* in sugar beet seedlings

Seedlings were collected over five time points (Table 5) for DNA extraction and qPCR to validate that *A. cochlioides* DNA could be detected in the seedling tissue, and to determine if *A. cochlioides* could be detected prior to development of visible symptoms. For naturally infested field soil, *A. cochlioides* DNA was first detected 5 DAP with a mean Ct value of 29.62, and consistently detected in all samples 7 DAP with a mean Ct of 25.03. For the used bioassay soil, *A. cochlioides* DNA was first consistently detected at 5 DAP, with a mean Ct value of 24.21. *A. cochlioides* DNA was consistently detected 2 days prior to visible ARR symptoms in both soil samples. These results confirmed that the qPCR assay could detect *A. cochlioides* in infected sugar beet seedling tissues ahead any visible symptoms.

**Table 5.** The qPCR-based detection of *Aphanomyces cochlioides* DNA in susceptible sugar beet seedlings planted in soil naturally infested with *A. cochlioides* and used bioassay soil. Ct values were derived from the fit point analysis..

| DAP<sup>a</sup> | Field soil naturally infested with *A. cochlioides* |  |  |  | Used bioassay soil |  |  |  |
| --- | --- | --- | --- | --- | --- | --- | --- | --- |
|  | Mean Ct value | No. of positive technical replicates | Technical replicates positive (%) | ARR symptoms present | Mean Ct value | No. of positive technical replicates | Technical replicates positive (%) | ARR symptoms present |
| 3 | NA<sup>b</sup> | 0 | 0 | No | NA | 0 | 0 | No |
| 5 | 29.62 | 3 | 75 | No | 24.21 | 4 | 100 | No |
| 7 | 25.03 | 4 | 100 | No | 20.97 | 4 | 100 | Yes |
| 9 | 23.87 | 4 | 100 | Yes | 21.87 | 4 | 100 | Yes |
| 11 | 17.27 | 4 | 100 | Yes | 17.56 | 4 | 100 | Yes |

<sup>a</sup> DAP = Days after planting
<sup>b</sup> NA = No amplification

### Detection of *A. cochlioides* in naturally infested soils

The qPCR assay was validated with DNA extracted from twelve soil samples that were naturally infested with *A. cochlioides*. The ARR bioassay confirmed each soil sample was infested with *A. cochlioides*, and determined the ARR DSI value. The ARR DSI values ranged from 48-100, with a mean DSI value of 87 (Table 6). Eleven samples tested positive for *A. cochlioides* DNA in every technical replicate, while one soil sample (Perley) was positive in 3 of 4 technical replicates. For 12 naturally infested soil samples the mean Ct values ranged from 26.72 - 34.64, indicating the assays capability in detecting *A. cochlioides* (Table 6).

**Table 6.**
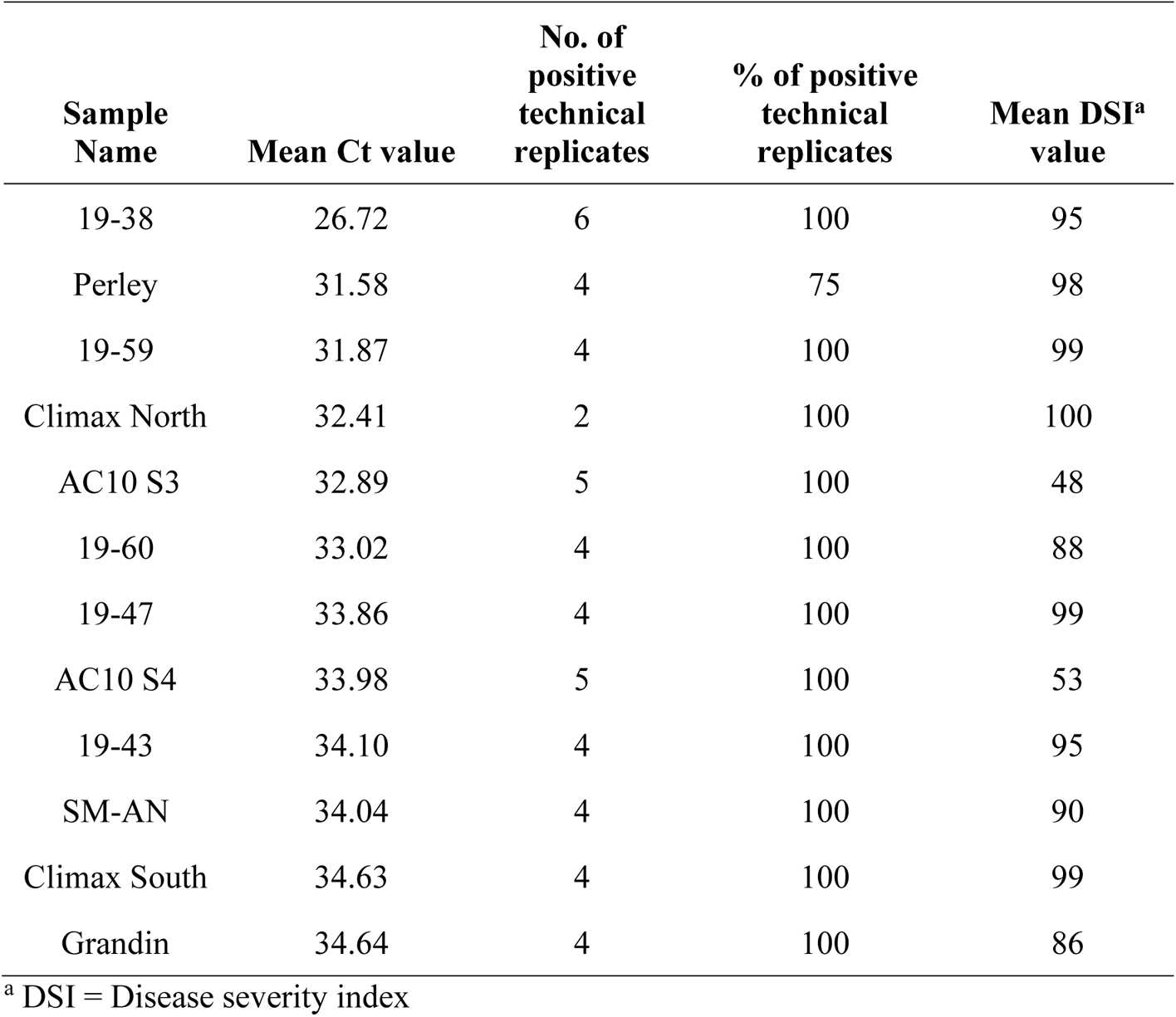
Ct values and Disease severity index (DSI) values for 12 field soil samples naturally infested with *Aphanomyces cochlioides*. Ct values were obtained using fit point analysis. Technical replicates with no qPCR amplification were not included in the calculation of mean Ct values.

### Detection of *A. cochlioides* in naturally infected plants

Out of 40 roots with ARR symptoms, culture-based diagnostics identified *A. cochlioides* in 23% of samples and *A. cochlioides* DNA was detected in 95% of samples. *A. cochlioides* DNA was detected in diseased sugar beet roots from every field whereas culture-based diagnostics identified *A. cochlioides* in 3 of 11 fields (Table 7). Finally, *A. cochlioides* DNA was detected in each of the 9 sugar beet roots that *A. cochlioides* was detected with using the culture-based assay, and in an additional 29 sugar beet roots where *A. cochlioides* was not recovered. These results validate that the qPCR assay can detect *A. cochlioides* in naturally infected mature sugar beet roots with high accuracy.

**Table 7.** Number of roots that *Aphanomyces cochlioides* was detected in using the qPCR and culture-based assay.

| Sample Name | DNA detection | Culture-based detection |
| --- | --- | --- |
| 20-51 | 4 | 0 |
| 20-52 | 4 | 0 |
| 20-53 | 7 | 3 |
| 20-54 | 2 | 0 |
| 20-55 | 2 | 0 |
| 20-56 | 1 | 0 |
| 20-57 | 4 | 0 |
| 20-58 | 3 | 1 |
| 20-59 | 2 | 0 |
| 20-60 | 7 | 5 |
| 20-61 | 2 | 0 |
| Total | 38 | 9 |

### Detection of *A. cochlioides* in artificially infested potting soil

The qPCR assay and ARR bioassay were performed on infested potting soil samples with various densities of *A. cochlioides* oospores to determine their mean Ct values and ARR DSI value and compare the two assays. In this experiment, the LOD for the qPCR assay was 10 oospores/g soil (dry wt.), which was consistently detected in all technical replicates, and had a mean Ct value of 33.26 (Table 8). A strong negative linear correlation (*R^2^*= 0.96, *p* = 0.00054) was observed between oospore density and mean Ct value. *A. cochlioides* DNA was not detected in the negative control, or in the lowest oospore density of 1 oospore/g soil (dry wt.). After completing the ARR bioassay for the soils, the mean ARR DSI value was calculated for each oospore density. The mean ARR DSI values ranged from 10.83 for 1 oospores/g soil (dry wt.) to 75 for 100 oospores/g soil (dry wt.) (Table 8). A strong positive linear correlation (*R^2^*= 0.968, *p* = 0.000032) was observed between the oospore density of the soil and the mean ARR DSI value. These results showed that the qPCR assay can detect *A. cochlioides* DNA in oospore infested potting soil, which had a range of DSI values from 23.33 to 75.00.

**Table 8.**
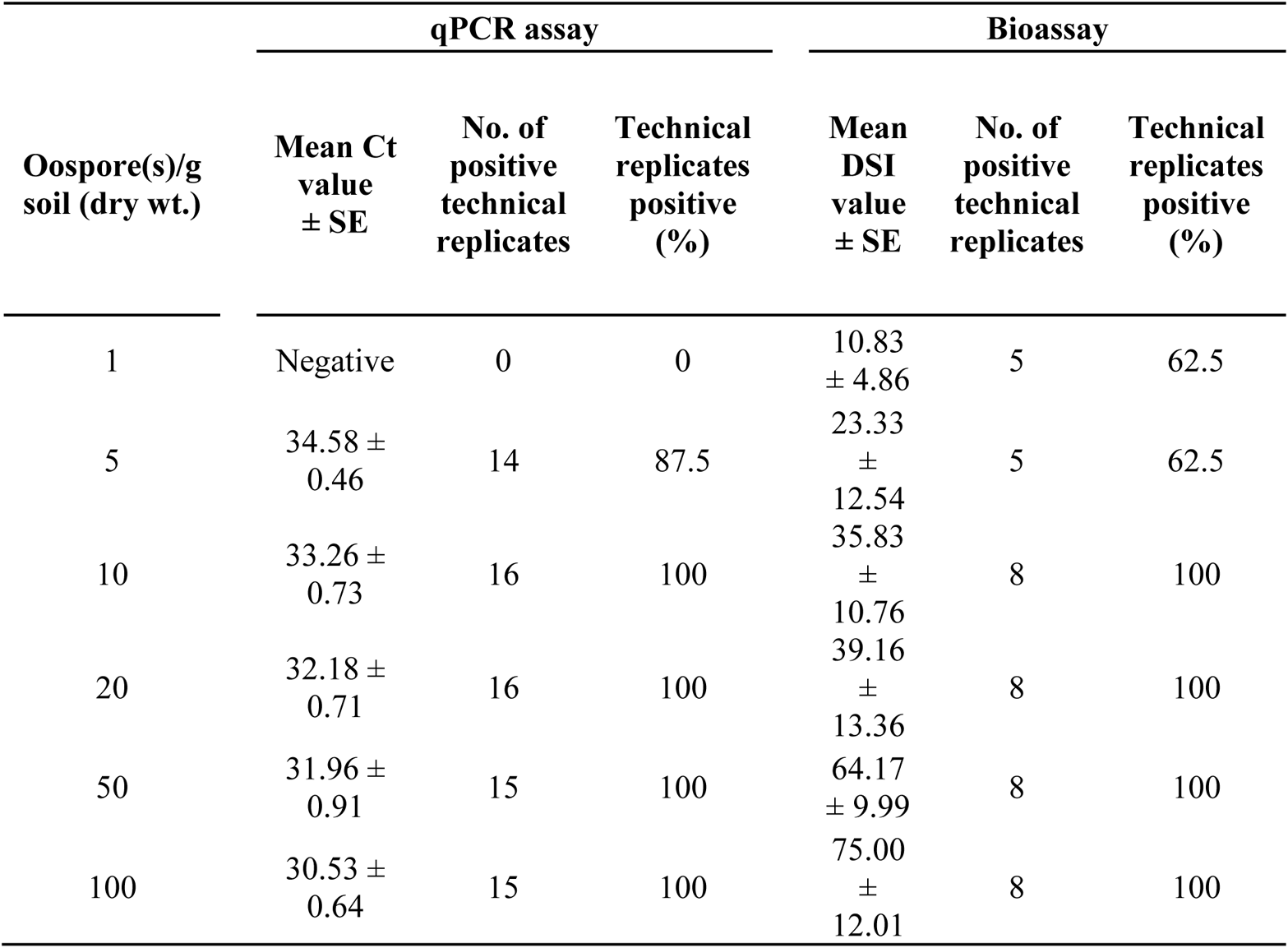
Mean Ct values and Aphanomyces root rot disease severity index (DSI) values for *Aphanomyces cochlioides* oospore infested potting soil samples.

| Oospore(s)/g<br>soil (dry wt.) | qPCR assay |  |  | Bioassay |  |  |
| --- | --- | --- | --- | --- | --- | --- |
|  | Mean Ct<br>value<br>± SE | No. of<br>positive<br>technical<br>replicates | Technical<br>replicates<br>positive<br>(%) | Mean<br>DSI<br>value<br>± SE | No. of<br>positive<br>technical<br>replicates | Technical<br>replicates<br>positive<br>(%) |
| 1 | Negative | 0 | 0 | 10.83<br>± 4.86 | 5 | 62.5 |
| 5 | 34.58 ±<br>0.46 | 14 | 87.5 | 23.33<br>±<br>12.54 | 5 | 62.5 |
| 10 | 33.26 ±<br>0.73 | 16 | 100 | 35.83<br>±<br>10.76 | 8 | 100 |
| 20 | 32.18 ±<br>0.71 | 16 | 100 | 39.16<br>±<br>13.36 | 8 | 100 |
| 50 | 31.96 ±<br>0.91 | 15 | 100 | 64.17<br>± 9.99 | 8 | 100 |
| 100 | 30.53 ±<br>0.64 | 15 | 100 | 75.00<br>±<br>12.01 | 8 | 100 |

## Discussion

The ultimate goal of developing qPCR assays for soilborne pathogens is qualitative detection and quantification of pathogen inoculum in field soils in order to assess the risk level of disease prior to planting. Having this information enables growers to make informed management decisions concerning which crop to plant, choosing varieties with appropriate level of disease resistance, use and rates of seed treatment, and other cultural management strategies. We report development of a specific, sensitive, and rapid mitochondrial DNA-based qPCR-based detection assay for *A. cochlioides* from various soil and plant samples. The assay was species-specific when tested against other *Aphanomyces* spp. and common soilborne fungi and oomycetes and detected all 14 isolates of *A. cochlioides* collected from MN, ND and TX. The gene order unique to *Aphanomyces* that was targeted improves specificity and repeatability across different real time PCR thermal cyclers. The forward primer (Aphcoc-F) was designed to anneal to multiple *Aphanomyces* spp., including pathogens of both plants and aquatic fish. The reverse primer (Aphcoc-R) was designed to anneal to *Aphanomyces* spp. that are plant pathogens; including *A. cladogamus, A. cochlioides*, and *A. euteiches*. The probe (Aphcoc-Pr) was designed to anneal to *A. cochlioides*. Based on alignments of this locus (Figure S1) modifying bases in the probe sequence would enable the development of a qPCR assay that can targets the mitochondrial genome of *A. cladogamus*, or *A. euteiches* while using the same forward and reverse primers developed in this study. The mitochondrial genome was chosen as the amplification target because of its high copy number, which may improve sensitivity. Also, mitochondrial genomes have been found to evolve more rapidly compared to nuclear genomes, making it easier to identify variation among closely related species for primer and probe design (Makkonen et al., 2016). The standard curve of *A. cochlioides* DNA isolated from pure cultures indicated that as little as 1 fg of pathogen DNA could be accurately detected with this assay. The LOD for oospore infested potting mix was 10 oospores per gram (dry wt.) of soil, which highlights the sensitivity of this assay. Although, the DNA standards for the qPCR assay were not run in the background of soil or plant DNA, therefore, the LOD of 1 fg does not necessarily represent the sensitivity that would be observed for some real-world samples, such as soil DNA that contains PCR inhibitors. For soil DNA samples from the field there is an expected reduction in template DNA amplification, compared to the PCR amplification of the same quantity of *A. cochlioides* DNA obtained from pure culture. For example, when 0.1 pg of *A. cochlioides* DNA was added to various soil DNA samples the Ct values were 0.69-2.14 higher than that from testing 0.1 pg of *A. cochlioides* DNA without soil DNA in the background (details of the qPCR inhibition are summarized in the Supplementary Text and Supplementary Table S1). In order to determine the actual sensitivity of this assay for naturally infested soil or infected plant samples from a specific field, DNA standards need to be tested with the appropriate background sugar beet or soil DNA to account for PCR inhibitors.

Different modifications have been made to DNA extraction procedures in order to improve the sensitivity of this qPCR assay. During the development of a qPCR assay for *V. dahliae*, improved detection was achieved by pre-treating samples with three cycles of freezing in liquid nitrogen for 2 min followed by heating to 70 ℃ (Bilodeau et al., 2012). In Almquist et al., (2016), they tested heating the soil samples and lysis buffer for 10 min at 65 ℃, and the addition of skim milk powder prior to DNA extraction, but neither technique improved the LOD for *A. cochlioides* DNA. In this study, freezing and heating cycles, as in Bilodeau et al., (2012), were applied to soil samples naturally infested with *A. cochlioides*, and detection improved for some samples, but lacked consistency (data not shown).

During periods of dormancy, the oospores of *A. cochlioides* can persist in field soil for a decade (Papavizas and Ayers, 1974). These durable thick-walled oospores within soil samples present a challenge for DNA extractions, and therefore detection of *A. cochlioides* in naturally infested soil samples. Various methods and DNA extraction kits have been employed in attempts to determine reliable techniques to acquire DNA from these samples for qPCR diagnostic assays. We utilized bead beating, which has been shown to be a dependable method of mechanical cell lysis for extracting DNA from soil samples infested with *A. cochlioides* and *A. euteiches* oospores (Almquist et al., 2016; Sauvage et al., 2007).

Previous research showed that the FastDNA Spin Kit for Soil with a sample size of 350 mg was preferred over the DNeasy PowerMax Soil Kit with a sample size of 10 g because the larger sample size did not improve the LOD, was more time consuming, and more costly for detecting *A. cochlioides* (Almquist et al., 2016). We determined the FastDNA Spin Kit for Soil with a sample size of 500 mg was preferable to the DNeasy PowerMax Soil Kit with a sample size of 5 g. We found the FastDNA Spin Kit for Soil had a higher DNA extraction efficiency, produced more overall positive samples, and resulted in lower Ct values for soils with active *A. cochlioides* (data not shown). Therefore, the FastDNA Spin Kit for Soil was chosen for conducting the soil DNA extractions in this study.

When DNA is extracted from soil samples, PCR inhibitors like humic acid can reduce or inhibit DNA amplification during qPCR (Hebda & Foran, 2015). Field soil, which was confirmed to be positive for *A. cochlioides* using the bioassay, tested negative for *A. cochlioides* DNA when the standard protocol of the FastDNA Spin Kit for Soil was used. Before modifications were made to the DNA extraction protocol, this field soil DNA was tested for qPCR inhibition using the spiked positive technique, where 0.1 pg *A. cochlioides* DNA was added to each soil DNA sample. The results showed that each soil DNA sample had slightly reduced amplification, compared to the amplification of 0.1 pg DNA without the background soil DNA, but no reaction was completely inhibited (Table S1). Some qPCR assays for field soil DNA samples have successfully utilized internal controls to determine when a sample has potential inhibition, and provide more accurate quantification results (Bilodeau et al., 2012; Burkhardt et al. 2018). In the future, the addition of an internal control multiplexed with the *A. cochlioides* assay would reduce the number of assays that need to be run.

Several existing qPCR assays for soilborne plant pathogens have successfully utilized the FastDNA Spin Kit for Soil and FastPrep instrument for DNA extractions, but had varied durations and speed for mechanical cell lysis (Almquist et al., 2016; Bilodeau et al., 2012; Wallenhammar et al., 2012). In qPCR assays for *A. cochlioides* and *P. brassicae*, the cell lysis duration was 30 s at a speed of 5.5 m/s (Almquist et al., 2016; Wallenhammar et al., 2012). In Bilodeau et al., (2012), the cell lysis duration was 45 s at a speed of 6.5 m/s, which successfully extracted *V. dahliae* DNA from microsclerotia in the soil. In our study, a cell lysis duration of 40 s at speed 6.0 m/s consistently extracted *A. cochlioides* DNA from infected sugar beet tissue and bioassay soil where *A. cochlioides* was active. However, this lysis duration and speed was unsuccessful at extracting *A. cochlioides* DNA from naturally infested field soil, where *A. cochlioides* was likely in a dormant oospore state. Increasing the cell lysis time and speed to 80 s and 7.5 m/s, respectively, with the addition of an extra ceramic bead allowed for the detection of *A. cochlioides* DNA from naturally infested soil samples with this qPCR assay. The optimization of the DNA extraction resulted in more positive samples and lower Ct values for each sample type we tested, therefore improving the detection of *A. cochlioides*.

Under field conditions, symptoms of ARR can be confounding, especially if fields dry out after initial infections which results in only superficial scarring on the surface of roots without any active rotting. We have observed that roots with scarring have poor rates of *A. cochlioides* recovery in the culture-based assay. It is also common for diseased roots to have mixed infections with other root rot fungi such as *Rhizoctonia solani* or *Fusarium* spp., which can outgrow *A. cochlioides* in culture and compromise its recovery. This qPCR assay reliably detected *A. cochlioides* in 95% of adult roots with ARR symptoms, compared to culture-based recovery where only 23% of the same roots were positive. Each of the roots that were positive with the culture-based assay were also positive with the qPCR assay, and the qPCR assay detected *A. cochlioides* DNA in an additional 29 other sugar beet roots where the pathogen was not cultured. This suggests that *A. cochlioides* DNA can be detected in adult diseased sugar beets after the pathogen is no longer active. Having this information expands our ability to detect this pathogen, and provides valuable information to growers concerning the presence of ARR in their fields. *A. cochlioides* causes post-emergence damping-off of seedlings, and it can be challenging to distinguish damping-off caused by other pathogens such as *R. solani* and *Pythium* spp. under field conditions. Infected seedlings tend to have higher rates of *A. cochlioides* recovery in the culture-based assay, but the presence of *R. solani* or *Pythium* spp. can compromise the recovery of *A. cochlioides*. This qPCR assay detected *A. cochlioides* in seedlings as early as 5 DAP and before the appearance of visible damping-off symptoms at 7 to 9 DAP. The ease of *A. cochlioides* detection in infected plant material highlights the value of using a culture-independent assay.

The qPCR assay developed in this study was validated using naturally infested field soil and oospore infested potting soil, naturally infected adult sugar beet roots, and sugar beet seedlings. Recently, a qPCR assay was developed to quantify *A. cochlioides* DNA from soil samples with primers that target a rRNA gene in the nuclear genome (Almquist et al., 2016). This qPCR assay was specific to *A. cochlioides*, and had an LOD in the range of 1-50 oospores/g soil (dry wt.) depending on soil characteristics, which was similar to qPCR assays for other oomycetes (Almquist et al., 2016). They found their assay was more sensitive for soils with higher clay content. For artificially oospore infested soil, they observed a consistent LOD of 10 oospores/g soil (dry wt.), which was also observed in our study. The Almquist et al., (2016) qPCR assay was only tested on soil samples, and consistently detected *A. cochlioides* in naturally infested soil samples with a DSI above 75, detected *A. cochlioides* in 50% of the samples when the DSI was 64-74, and did not detect *A. cochlioides* when DSI was below 50. In comparison, for naturally infested soil samples, our qPCR assay detected *A. cochlioides* when DSI values were as low as 48 (mean Ct = 32.89), although the majority of these samples (10 out of 12) had DSI values above 86. Furthermore, for oospore infested potting soil our qPCR assay detected *A. cochlioides* when mean DSI values ranged from 23.33-75. This assay had a high correlation (*R^2^*= 0.96) with oospore inoculum density ranging from 5 to 100 oospores per gram of soil using an artificially infested potting mix. However, for the bioassay of the artificially infested potting mix there was significant variability between the technical replicates, so means were used to compare the oospore density and soil DSI value (Table 8). Quantification of *A. cochlioides* at these levels is very useful for sugar beet growers to devise a disease management plan. In the future, validating these results using field soil with a broader range of DSI values would support the reliability of this assay for accurate quantification. Overall, this qPCR assay may be a useful tool to provide accurate information to sugar beet growers concerning the quantity of *A. cochlioides* inoculum in their fields.

Quantitative PCR assays for soilborne pathogens aim to provide an accurate estimate of disease potential in the soil. While this assay has been demonstrated to be effective for pathogen detection with asymptomatic seedlings, adult plants exhibiting root rot symptoms, and from infested soil samples, before it can be effectively used for quantification of the pathogen in field soil samples additional work is needed. This would include 1) multiplexing with an internal control to improve evaluation of PCR inhibition while reducing the number of assays that need to be run; 2) rerunning the standard curve with background DNA extracted from pathogen-free soil or plant samples to evaluate the effectiveness of DNA extraction techniques and provide an LOD that accurately reflects conditions under which the assay will be used; 3) evaluating field sampling strategies to determine the number of samples needed to accurately evaluate risk of ARR for the grower. The importance of this was highlighted by the variable qPCR results observed for two soil subsamples from the same infested field soil (data not shown). While it could be due to presence of oospore inoculum in disintegrating organic matter, additional experimentation is needed to reduce this level of variation; and 4) determine how the ARR DSI of a field soil correlates with the *A. cochlioides* Ct value for naturally infested soils. It will be vital to apply an optimal sampling strategy to provide an accurate representation of risk for a field, and test a large collection of soil samples with a wide range of DSI values. It would also be useful to evaluate the effect of soil type and microflora on the predicted disease potential compared to the realized diseased in the field.

## Acknowledgements

We would like to thank American Crystal Sugar Company, Minn-Dak Farmers Cooperative, Southern Minnesota Beet Sugar Cooperative, and USDA-NIFA for funding our research.

## Literature Cited

Agrios, G. M. 2005. Plant Pathology. p. 391. Elsevier Inc. Burlington, MA.

Alexiades, A., Kendall, A., Winans, K. S., and Kaffka, S. R. 2018. Sugar beet ethanol *(Beta vulgaris* L.): A promising low-carbon pathway for ethanol production in California. J. Clean. Prod. 172:3907–3917.

Almquist, C., Persson, L., Olsson, Å., Sundström, J., and Jonsson, A. 2016. Disease risk assessment of sugar beet root rot using quantitative real-time PCR analysis of *Aphanomyces cochlioides* in naturally infested soil samples. Eur. J. Plant Pathol. 145:731–742.

American Crystal Sugar Company (ACSC). n.d.. Sugar Processing. Available from https://www.crystalsugar/processing.com. Accessed 19 September 2020.

Beale, J. W., Windels, C. E., and Kinkel, L. L. 2002. Spatial Distribution of *Aphanomyces cochlioides* and Root Rot in Sugar Beet Fields. Plant Dis. 86(5):547–551.

Bilodeau, G. J., Koike, S. T., Uribe, P., and Martin, F. N. 2012. Development of an assay for rapid detection and quantification of *Verticillium dahliae* in soil. Phytopathology. 102:331–343.

Burkhardt, A., Ramon, M. L., Smith, B., Koike, S.T. and Martin, F. N. 2018. Development of molecular methods to detect *Macrophomina phaseolina* from strawberry plants and soil. Phytopathology. 108:1386–1394

Dohm, J. C., Minoche, A. E., Holtgräwe, D., Capella-Gutiérrez, S., Zakrzewski, F., Tafer, H., Rupp, O., Sörensen, T. R., Stracke, R., Reinhardt, R., Goesmann, A., Kraft, T., Schulz, B., Stadler, P. F., Schmidt, T., Gabaldón, T., Lehrach, H., Weisshaar, B., and Himmelbauer, H. 2014. The genome of the recently domesticated crop plant sugar beet (*Beta vulgaris*). Nature. 505:546–549.

Draycott, A. P. 2006. Sugar Beet. Ames, Iowa: Wiley-Blackwell. pp. 286–293. Available from http://base.dnsgb.com.ua/files/book/Agriculture/Cultures/Sugar-Beet.pdf

Dyer, A. T., and Windels, C. E. 2003. Viability and maturation of *Aphanomyces cochlioides* oospores. Mycologia. 95:321–326.

Fink, H. C., and Buchholtz, W. F. 1954. Correlation between sugar beet crop losses and greenhouse determinations of soil infestation by *Aphanomyces cochlioides*. Am. Soc. Sugar Beet Technol. 8:252–259.

Grünwald, N. J., and Coyne, C. J. 2003. Species of Aphanomyces described as plant pathogens including known hosts and names of diseases. Page 13 in: Proceedings of the Second International Aphanomyces Workshop, U.S. Department of Agriculture, Agricultural Research Service, Pasco, Washington. Available from http://sites.science.oregonstate.edu/bpp/labs/grunwald/publications/ProceedingsAphanomycesWorkshop.pdf

Harveson, R. M. 2007. Aphanomyces Root Rot of Sugar Beet. NebGuide G1407. University of Nebraska-Lincoln Extension, Institute of Agriculture and Natural Resources. Available from http://extensionpublications.unl.edu/assets/pdf/g1407.pdf

Harveson, R. M., Windels, C. E., Smith, J. A., Brantner, J. R., Cattanach, A. W., Giles, J. F., Hubbell, L., and Cattanach N. R. 2007. Fungicide Registration and a Small Niche Market: A Case Study of Hymexazol Seed Treatment and the U.S. Sugar Beet Industry. Plant Dis. 91:780–790.

Hebda, L. M., and Foran, D. R. 2015. Assessing the Utility of Soil DNA Extraction Kits for Increasing DNA Yields and Eliminating PCR Inhibitors from Buried Skeletal Remains. Forensic Sci. 60:1322–1330.

Islam, M. T. 2010. Morphology and behavior of the successive generations of zoospores of a damping-off pathogen *Aphanomyces cochlioides*. Plant Pathol. 92:461–468.

Makkonen, J., Vesterbacka, A., Martin, F., Jussila, J., Diéguez-Uribeondo, J., Kortet, R., and Kokko, H. 2016. Mitochondrial genomes and comparative genomics of *Aphanomyces astaci* and *Aphanomyces invadans*. Sci. Rep. 6:36089.

Olsson, Å., Persson, L., and Olsson, S. 2011. Variations in soil characteristics affecting the occurrence of Aphanomyces root rot of sugar beet – Risk evaluation and disease control. Soil Biol. Biochem. 43:316–323.

Olsson, Å., Persson, L., and Olsson, S. 2019. Influence of soil characteristics on yield response to lime in sugar beet. Geoderma. 337:1208–1217.

Papavizas, G. C., and Ayers, W. A. 1974. Aphanomyces species and their root diseases in pea and sugarbeet. U. S. Dept. Agriculture, Agric. Res. Serv., Tech. Bull. 1485:158.

Sauvage, H., Moussart, A., Bois, F., Tivoli, B., Barray, S., and Laval, K. 2007. Development of a molecular method to detect and quantify *Aphanomyces euteiches* in soil. FEMS Microbiol. Lett. 273:64–69.

U. S. Department of Agriculture National Agricultural Statistics Service. 2022. National Statistics for Sugarbeets. Available from https://www.nass.usda.gov/Statistics_by_Subject. Accessed 7 April 2022.

Vandemark, G. J., Kraft, J. M., Larsen, R. C., Gritsenko, M. A., and Boge, W. L. 2000. A PCR-Based Assay by Sequence-Characterized DNA Markers for the Identification and Detection of *Aphanomyces euteiches*. Phytopathology. 90:1137–1144.

Wallenhammar, A. C., Almquist, C., Söderström, M., and Jonsson, A. 2012. In-field distribution of *Plasmodiophora brassicae* measured using quantitative real-time PCR. Plant Pathol. 61:16–28.

Weiland, J. J., and Shelver, W. L. 2004. Production and Characterization of Antiserum to *Aphanomyces cochlioides*. J. Sugar Beet Res. 41:179–190.

Weiland, J. J., and Sundsbak, J. L. 2000. Differentiation and Detection of Sugar Beet Fungal Pathogens Using PCR Amplification of Actin Coding Sequences and the ITS Region of the rRNA Gene. Plant Dis. 84:475–482.

Windels, C. E. 2000. Aphanomyces root rot on sugar beet. Plant Health Prog. https://doi.org/10.1094/PHP-2000-0720-01-DG

Windels, C. E., and Brantner, J. R. 2000. Variability of spore production and aggressiveness of *Aphanomyces cochlioides* on sugarbeet. Sugarbeet Research and Extension Reports. 31:241–246.

Windels, C. E., and Brantner, J. R. 2001. Benefit of Tachigaren - treated sugarbeet seed in soils with different Aphanomyces soil index values. Sugarbeet Research and Education Board. Available from https://www.sbreb.org/wp-content/uploads/2018/12/01-Benefit-of-Tach-Carol.pdf. Accessed 7 October 2020.

Windels, C. E., and Nabben-Schindler, D. J. 1996. Limitations of a greenhouse assay for determining potential of Aphanomyces root rot in sugarbeet fields. J. Sugar Beet Res. 33:1–4.

Wen, Y., Islam, M. T., and Tahara, S. 2006. Phenolic Constituents of *Celosia cristata* L. Susceptible to Spinach Root Rot Pathogen *Aphanomyces cochlioides*. Biosci., Biotech., Biochem. 70:2567–2570.

