## Supplementary for "DNA-based detection of *Aphanomyces cochlioides* in soil and sugar beet plants"

**Supplementary Information**

**Evaluation of qPCR inhibition in soil DNA samples**. Prior to optimizing the FastDNA Spin Kit for Soil extraction procedure, the spiked positive technique was used to evaluate qPCR inhibition for soil samples that tested negative for *A. cochlioides* DNA using the qPCR assay. This technique also revealed how amplification of *A. cochlioides* DNA is altered when in the presence of various DNA from soil. The soil DNA samples were spiked with 0.1 pg of pure *A. cochlioides* DNA. In addition, the 0.1 pg DNA standard was tested by itself in the same qPCR plate, which produced a mean Ct of 26.66. In the absence of PCR inhibition, the Ct values of each soil DNA sample with the spiked 0.1 pg DNA will be 26.66 or lower. In the presence of any inhibition, the Ct values of the soil sample spiked with 0.1 pg DNA will be higher than 26.66. If there is no qPCR amplification for the spiked soil DNA sample, while the 0.1 pg DNA standard alone produces a normal Ct value, this signifies complete inhibition of PCR.

Prior to the optimization of the DNA extraction, the qPCR assay was not able to detect *A. cochlioides* in DNA samples from soils confirmed to be naturally infested with *A. cochlioides* using the soil bioassay. This soil DNA was extracted using the standard protocol for the FastDNA Spin Kit for Soil, and was negative for *A. cochlioides* DNA after being tested with the qPCR assay. Due to the concern that inhibition may have been causing these samples to be negative, the spiked positive technique was used to evaluate inhibition during the qPCR amplification of these samples. The Ct difference was calculated by taking the mean Ct value of each sample and subtracting the mean Ct value of the 0.1 pg positive control (26.65). Across all soil DNA samples, the range of Ct difference was 0.69-2.14, and the mean Ct difference was 1.25 (Table S1). The results showed that the PCR amplification of 0.1 pg of pure *A. cochlioides* DNA was reduced slightly in the presence of various soil DNA, but never inhibited.

**Supplementary Table S1**. Mean Ct values of naturally infested soil sample DNA extracted with the standard protocol of the FastDNA Spin Kit for Soil with spiked positive control (PC). The PC was 0.1 pg *Aphanomyces cochlioides* DNA with a consistent Ct value of 26.65.

| **Sample Name** | **Mean Ct value** | **Ct value deviation from PC** |
| --- | --- | --- |
| AC10 S3 + PC | 27.79 | 1.14 |
| AC10 S4 + PC | 27.34 | 0.69 |
| 19-38 + PC | 28.05 | 1.4 |
| 19-43 + PC | 28.79 | 2.14 |
| 19-47 + PC | 27.82 | 1.17 |
| 19-59 + PC | 27.99 | 1.34 |
| 19-60 + PC | 27.89 | 1.24 |
| SM-AN + PC | 27.99 | 1.34 |
| Grandin + PC | 27.49 | 0.84 |
| Climax South + PC | 27.55 | 0.9 |
| Climax North + PC | 28.36 | 1.71 |
| Perley + PC | 27.72 | 1.07 |

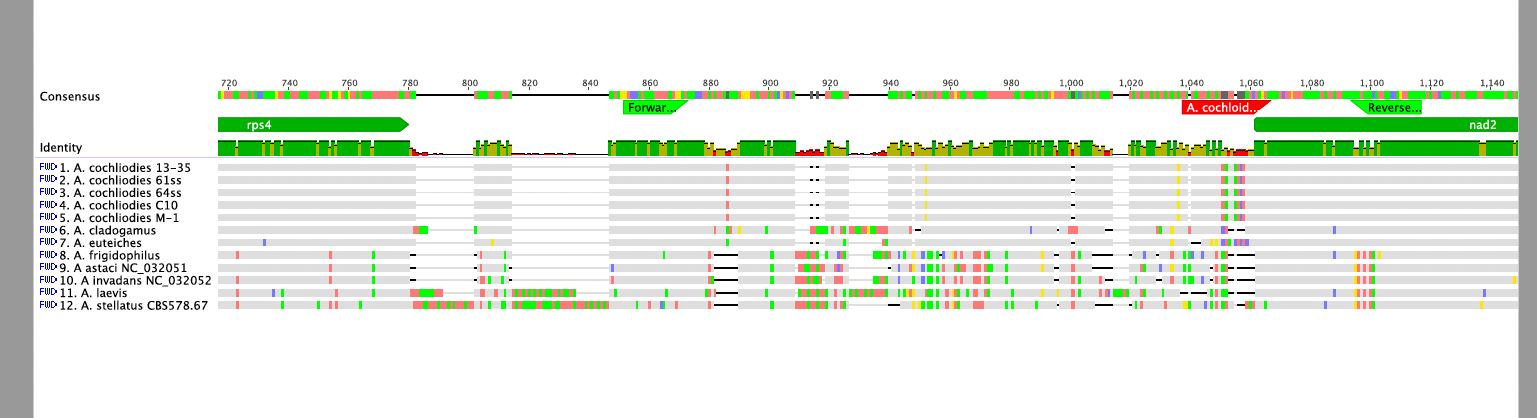

**Supplementary Figure S1**. Specificity of Aphcoc-F (forward) and Aphcoc-R (reverse) primers (light green), and probe Aphcoc-Pr (red) amongst 5 isolates of *A. cochlioides* and other closely related *Aphanomyces* spp*.*.
